# Using regularized regression and biological covariation to impute missing values in quantitative proteomics

**DOI:** 10.1101/2025.10.27.684979

**Authors:** Matthew Sonnett, Leonid Peshkin, Marc W Kirschner

**Affiliations:** Department of Systems Biology, Harvard Medical School, Boston, MA, USA, 02115

## Abstract

Proteomics studies analyzing many samples typically generate datasets with missing values where many protein abundances are only quantified in a subset of assayed conditions. While multiplexing with isobaric tags can address this by combining multiple samples into a single injection, missing values are unavoidable when the sample count exceeds the number of available isobaric tags (currently >35). Such missing data complicates the interpretation of large-scale studies across diverse experimental conditions. Here, we introduce a method to impute missing values of relative protein abundance by leveraging measurements from other proteins in the dataset through regularized regression. Our technique, which is applicable to diverse datasets including different cell lines, animals, or biochemical perturbations, capitalizes on the hitherto overlooked biological covariation among protein abundance changes. Our analysis of eight published proteomics datasets reveals a robust imputation capability, achieving a median R^2^ of 0.55 to 0.8 between imputed and measured data. We demonstrate a similar imputation efficacy in multiple measurement modalities: TMT, DIA, label free, and TMT phosphoproteomics. When examining regression coefficients that were pivotal for accurate data imputation we found that those often mirror known biology. We propose that previously overlooked biological covariation might lead to the generation of novel hypotheses and ultimately advance our understanding of systems level protein organization.

## Introduction

The predominant method in contemporary proteomics is peptide fragment identification via mass spectrometry (MS). However, MS-based peptide identification is inherently biased and stochastic. The preferential analysis of higher abundance species and the variable coverage in sequential MS analyses result in inconsistent peptide identification even in technical replicates (Fig S1A). While mapping peptides to single protein measurements alleviates some of the incompleteness, missing values remain an issue, especially for proteins identified by a minimal number of peptides (Fig S1B).

As biological experiments grow in complexity and size the issue of incomplete data magnifies. For example, a study of the protein composition in different immune cell types has a union of 10,012 identified proteins among 97 samples, but only 2,390 proteins were identified and measured in all 97 samples (Fig S2A) (Rieckmann, 2017). This problem can only be partially remedied using multiplexed tags. Current multiplexing approaches permit up to 35 samples to be barcoded and combined into a single run (Thompson, 2019 and Zuniga, 2024). More than a single run of TMT multiplexed samples however can be co-analyzed by using a common sample in different experiments as a “bridge” (Lapek, 2017). However, as the results from different TMT experiments are aggregated, there is still significant loss of data (Fig 1A). For example, in a large-scale TMT study of 378 cancel cell lines (Nusinow, 2020), a total of 12,755 proteins were quantified in at least one cell line, but only 5,153 proteins were quantified in all 378 (Fig 1A, orange). These commonly identified proteins were, as expected, the most abundant. However, there was a noticeable deficiency of low abundance proteins of interest such as transcription factors and signaling molecules in the combined dataset. Out of the 765 transcription factors quantified in at least one cell line, only 106 were quantified across all (Fig S2B). The challenge further intensifies in state-of-the-art single cell experiments, evidenced by a study which, despite identifying 2,844 proteins across 1,543 profiles, had consistent identification for only 13 proteins (Fig S3) (Leduc, 2022).

**Figure 1.**
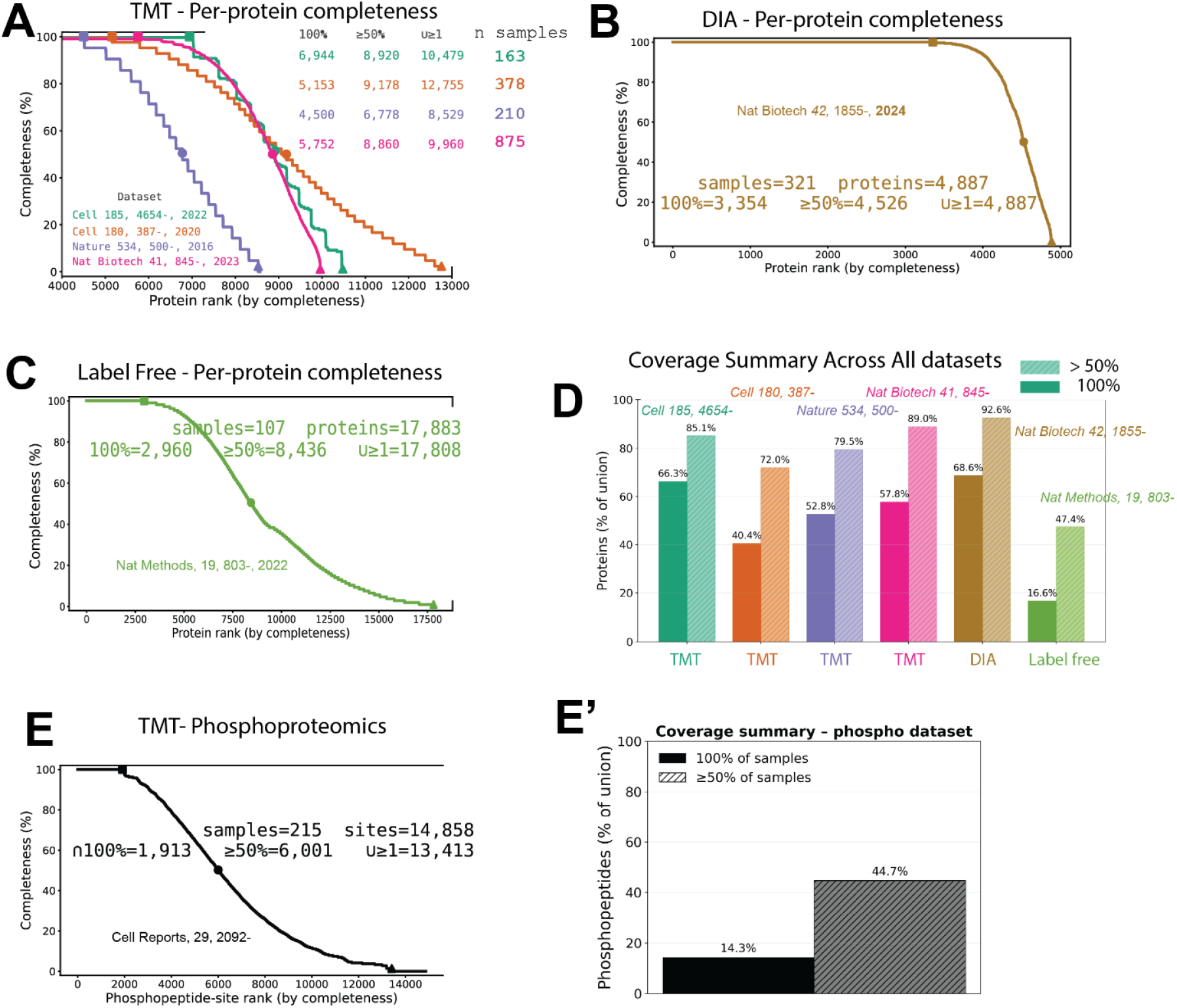
The missing value problem in proteomics across methods. **A)** TMT Protein level. Per protein completeness curves for several representative TMT datasets. Proteins are ranked by completeness (y-axis). Annotated numbers indicate the total number of samples (n), the number of proteins observed in 100% of samples, the number observed in ≥ 50% of samples, and the union (u ≥ 1). As the number of samples analyzed increases, the set of proteins quantified in all samples decreases resulting in missing values. **B)** DIA Protein level. Completeness curve for a representative DIA study in yeast. Despite a smaller genome, only 68% of all proteins observed are quantified in every sample. **C)** Label free DDA Protein level. Completeness curve a large label free dataset of different mouse organs. A comparatively larger gap between the union and 100% set is observed. **D)** Cross study coverage summary: bars show, as a percentage of the union within each dataset, the fraction of proteins quantified in 100% of samples (solid), and >50% of samples (hatched). Colors correspond to the same studies as in A-C. Across all studies and proteomics methods, the 100% set is consistently much smaller than the union. **E)** Completeness curve for TMT phosphopeptide site quantification in yeast. **E’)** Coverage summary of TMT yeast phophopeptide site data from E). Only 14.3% of phosphopeptide sites were measured in all 215 samples, and only 44.7% of sites were measured in ≥ 50% of samples.

Data independent acquisition (DIA) offers a distinct approach to address the challenge of missing values in proteomics. Contrary to the inherent unpredictability in which MS2 spectra are captured in traditional methods, DIA operates on a set acquisition schedule for MS2 spectra. Despite this systematic acquisition strategy, DIA still grapples with the presence of missing values. For example, a recent DIA study in yeast (Guzman, 2024) quantified a total of 4,887 yeast proteins, but only 3,354 were quantified in all 321 samples analyzed (Fig 1B). This is an improvement, but not a drastic one from an older TMT yeast study (Li, 2019), which quantified a total of 4,647 yeast proteins, with 3,369 quantified in all 215 samples analyzed (Fig S4). Furthermore, when benchmarked against experiments employing pre-fractionated samples paired with isobaric tags, DIA tends to suffer in terms of measurement precision and the overall proteomic coverage. Presently, for comparative analyses involving up to 35 conditions, isobaric tagging demonstrates superior data quality. However, for broader studies, the advantage is commonly thought to tilt towards DIA, primarily due to its reduced occurrence of missing values. While there are multiple advantages of DIA, particularly in some sample types such as those from blood, the missing value problem is still important and prevalent.

For these reasons it has long been a goal in quantitative proteomics to impute missing values. Multiple approaches have been tried with label free data, with varying degrees of success (Webb-Robertson, 2015). Various machine learning approaches including matrix decomposition (PCA), clustering (k-nearest neighbors) and linear discriminant analysis have been implemented (Webb-Robertson, 2015). These approaches have had limited success. One possible explanation for these difficulties is the use of too few samples. Another is the imprecision of the data (Sonnett, 2018). A popular approach is to sample a noise model such as a low valued gaussian with the assumption that an undetected peptide is likely to have a low value (Rieckman, 2017). However, without any inherent correlation between the imputed and actual values, any form of mean imputation in this context is probably an unsound approach (O’Brien, 2018). The “Match Between Runs” technique has made significant progress for its unique approach: rather than inferring values statistically, it directly locates peaks in the MS1 raw spectra by coupling high accuracy mass measurements and aligned retention times. Yet, even this approach hasn’t been a panacea, with its efficacy varying across different applications (Lim, 2019).

Leveraging biological covariation for statistical inference and imputation is a well-established concept particularly in the realm of genomics. Connectivity maps of molecules, genes, and diseases have been successfully generated using gene-expression signatures (Golub, 2006). To economize on RNA-seq measurements, Golub et al created a set of 1,000 representative genes to predict the abundance of all ∼20,000 transcripts (Subramanian, 2017). Furthermore, there has been a recent focus on the imputation of missing values in single cell RNA seq (Hou, 2020). Different approaches in single cell transcriptomics have used low-rank matrix approximation (Linderman, 2022), subspace regression (Tran, 2022), and Bayesian factorization (Wen, 2022). These methods, along with others, have improved the coverage of single cell RNA-seq measurement studies. However, there are data issues for single cell transcriptomic analysis that distinguish these from proteomic analysis. A major advantage in single cell transcriptomic studies is that many thousands of conditions are measured (each cell constitutes a condition). This allows the generation of greater data and greater statistical power based on co-variation. Typically, hundreds of conditions are analyzed in a large-scale proteomics study (Chick, 2016 and Gyuricza 2022).

Inspired by transcriptomic studies, we hypothesized that for different samples with varying experimental designs (*eg*. cell types, individual animals, drugs) extensive biological covariation between proteins in each sample may exists of sufficient power to enable us to infer the abundance of many proteins not previously measured. The most familiar example of this would be positive covariation that is observed between different protein members of the same protein-protein complex (Bludau, 2020). Indeed, not surprisingly, strong correlations between individual members of the protein complexes have been observed in multiple data sets (Lapek, 2017, Chick, 2016, Pan, 2018). Unfortunately, only a small number of proteins are known members of stable stoichiometric complexes. Our much more ambitious goal is to employ covariance without having to rely on any preexisting biological information for co-variation.

In our attempt to infer missing values, our foundational premise is that any protein within the dataset could potentially influence the variation of another. This entails constructing an expansive set of linear models. However, a challenge arises when the number of predictors (proteins, in this context) greatly surpasses the number of observations (distinct conditions). Such models face two primary issues. Firstly, the sheer volume of predictors leads to an overdetermined system devoid of a singular solution. Secondly, conventional least squares regression tends to assign non-zero coefficients to every protein, mistakenly implying each protein’s significance to the protein under examination. This is very likely to be wrong because in virtually all circumstances a *priori* simple explanations are thought to be much more likely to be true than extremely complex ones (Occam’s razor). This concept fits what we know about how many biological systems are built, where complex models of hundreds to thousands of proteins are not needed to understand most known biological mechanisms. Furthermore, there is a cost to adding large numbers of weak, noisy measurements, as the accumulated noise in weak correlations will inevitably diminish prediction accuracy. On the other hand, some biological processes may involve many components, and overly simple models may not provide sufficient predictive power.

Here, rather than predetermining covariation patterns, we allow the data itself to determine the most fitting complexity for each protein. To address the intricate challenge of model selection, we employ the elastic net technique (Zou, 2005). As a generalized form of regularized regression, the elastic net introduces constraints on the number of protein species considered, varying sparsity, while also managing the influence each protein holds, ensuring shrinkage. By exploring a myriad of potential models, our approach can select the best-suited model, whether it’s a simplistic one with just a few proteins or a comprehensive one that includes thousands. This methodology ensures that our chosen model strikes an optimal balance between sparsity and shrinkage, providing robust predictions without overfitting.

In this work, we develop a data-driven imputation strategy that uses elastic-net regularized regression to leverage biological covariation across proteins, and we validate it on held out ground truth values created from complete subsets of published datasets. We show that the approach accurately recovers missing protein (and phosphopeptide) abundances across diverse study designs and platforms, including TMT, DIA, label-free, and TMT-phosphoproteomics, with median prediction R^2^ typically ∼0.55–0.8, while preserving global data structure in PCA and hierarchical clustering and substantially increasing the number of analyzable features. We quantify how performance scales with missingness and cohort size, and we demonstrate that learned coefficients often recapitulate known complexes and pathways while also nominating plausible new positive and negative relationships. Together, these results establish a general, modality agnostic framework that expands proteome coverage, especially for low abundance regulators, and yields interpretable covariation patterns that can guide biological hypothesis generation in large-scale proteomics.

## Materials and Methods

### Data preprocessing

All proteomic matrices were log transformed and then column standardized (zero mean, unit variance) prior to modeling. Non-finite entries (NaN, ±Inf) were treated as missing. Matrices used for paired comparisons (measured vs imputed) were intersected on identical sample and protein sets while preserving order.

### Regularized regression and out-of-fold evaluation

For each target protein, its standardized abundance was modeled from all other proteins using elastic net (primary) and ridge (baseline) in an out-of-fold design with K = 5 folds unless stated otherwise. Elastic net was fit with celer (ElasticNetCV for tuning followed by ElasticNet for refit) using l1 ratios {0.2, 0.5, 0.8, 1.0} and 60 alphas, selecting the 1-SE alpha at the best l1 ratio. We used fit_intercept=True, tol=1e-3, and an adaptive max_iter (200→400→800 on convergence warnings). When celer was unavailable, we used scikit-learn’s compatible estimators. Ridge used RidgeCV with gcv_mode=“svd” over 60 log-spaced alphas spanning 10^−4^ to 10^3^; an optional inner-CV ridge with a 1-SE rule exists but was off by default. Out-of-fold predictions were collected for every sample/feature to yield unbiased performance estimates.

### Controlled missingness experiment

Unless specified otherwise, we simulated sparsity by masking 20% of sample rows per target uniformly at random (one replicate), fitting models on the observed subset, and scoring predictions only on the masked entries. A mean of train baseline was computed in parallel for OOF and sweep settings.

### Permuted-null controls

Figures labeled “permuted-null” were generated with ridge on a shuffled response: for each target, the response vector was permuted within feature prior to ridge fitting, with alpha chosen by GCV as above. We used this ridge_shuf procedure for both OOF and the 20% missingness sweep (ridge_shuf_sweep) to obtain an empirical null distribution of predictability given the same design matrix.

### Performance metrics

For OOF and sweep settings we reported RMSE, MAE, R^2^ with a variance-zero guard, and Pearson r, computed only on finite truth–prediction pairs. Where parametric summaries of correlation distributions were shown, we applied Fisher’s z transform.

### Principal component analysis and subspace similarity

PCA was performed with scikit-learn PCA (full SVD) on column centered matrices (samples as observations). Variance spectra were compared across measured, imputed, and column shuffled counterparts. For visualization and alignment, loading sign indeterminacy was resolved by flipping the signs of imputed loadings to maximize column wise correlation with measured loadings. Global geometry preservation between measured and imputed spaces was quantified by Procrustes alignment (scipy.spatial.procrustes) on the first k PCs and by the RV coefficient computed from centered score cross product scatter matrices.

### Clustering and agreement

Hierarchical clustering of samples used Ward linkage on Euclidean distances (scipy.cluster.hierarchy with pairwise distances from scikit-learn). Dendrograms were cut at prespecified k to form hard partitions. Agreement between partitions derived from measured vs imputed matrices was quantified by Adjusted Rand Index and Normalized Mutual Information (sklearn.metrics.adjusted_rand_score, normalized_mutual_info_score).

### Neighborhood preservation and correlation structure tests

Local structure was assessed with trustworthiness (sklearn.manifold.trustworthiness), comparing neighborhoods in the original feature space to neighborhoods in the PCA space for measured and imputed matrices. To compare proteome wide covariation, we drew up to 2×10^6^ random protein pairs without replacement, computed Pearson r across samples in measured and imputed matrices, and contrasted these distributions to their column-shuffled nulls using the first Wasserstein distance (scipy.stats.wasserstein_distance). Where applicable we reported ECDFs or kernel smoothed PDFs for visualization only.

### Software

Python packages used include numpy 2.3.3; scipy 1.16.2; pandas 2.3.3; scikit-learn 1.7.2; celer 0.7.4; matplotlib 3.10.7; seaborn 0.13.2; joblib 1.5.2. Random seeds were fixed where relevant for reproducibility; parallelization details do not affect statistical procedures and are omitted.

## Results

### Pervasive, scale-dependent missingness across proteomics modalities

Large-scale proteomics studies suffer from a fundamental, scale dependent loss of across sample coverage that is evident across all commonly used proteomics approaches. We began by quantifying “completeness”: the fraction of samples in which each protein (or phosphosite) is quantified, and visualizing how the complete case set collapses as cohorts grow. In four representative TMT experiments where the number of samples analyzed ranged from 163 to 875 (Chick 2016, Nusinow 2020, Xiao 2022, Mitchell 2023), per protein completeness curves show a pronounced separation between the union of all quantified proteins and those present in every sample (Fig. 1A). In these TMT experiments, typically ∼70 – 85% of all proteins are measured in half of the conditions (≥50% threshold, hatched bars), whereas only ∼40% to ∼60% of proteins are measured in every condition (100% threshold, solid bars) (Fig. 1D). DIA improves per run depth but exhibits the same signature of sample to sample dropout: in a 321 sample yeast study, 4,887 proteins are detected in total, yet only 3,354 (68%) are present in all samples (Fig. 1B) (Guzman, 2024). Label free DDA shows an even larger gap between the union and the 100% intersection in a very sensitive multi organ mouse dataset underscoring that run level stochasticity and study scale jointly erode complete case coverage (Fig. 1C) (Giansanti 2022). In this dataset only 16.6% of all proteins were quantified in every sample.

The deficit is most severe for low abundance and post translational measurements. In a large TMT cell line compendium (Nusinow, 2020), the aggregate union contains 765 transcription factors but the complete case set retains only 106 (13.8%) (Fig S2B), reflecting a strong abundance bias relative to the full dataset where 40.4% of all proteins are quantified in every sample (Fig 1D, orange). Likewise, in a 215 sample TMT phosphoproteomics cohort, only 14.3% of phosphosites are quantified in every sample and 44.7% meet the ≥50% threshold (Fig. 1E–E′). These cross study summaries make clear that even when single run depth is high, integrating dozens to hundreds of samples yields matrices with extensive, structured missingness. Biological analysis of proteomics data such as PCA, clustering etc. typically rely on complete data matrices that would discard as much as half of all proteins measured.

These observations motivate the need for an imputation strategy that expands the analyzable matrix rather than restricting to complete cases. We model each protein from the rest of the proteome and predict its missing entries using out-of-fold regularized regression (elastic net as the primary model with ridge comparators), then retain predictions that meet a user-specified error criterion. This procedure yields matrices that are complete, or substantially enlarged, while preserving global structure (PCA, clustering). In practice, the analyzable feature set often increases by roughly two-fold across TMT, DIA, label free, and phosphoproteomics datasets, enabling downstream analyses that would otherwise discard large swaths of biologically informative proteins.

### Biological covariation enables accurate regression based imputation

Judging the effectiveness of a regularized regression approach first requires that we have a dataset where we know the ground truth and can thereby evaluate our predictions. Any existing data set of multiple experiments we would use would be expected to contain missing values, i.e. proteins appearing in one set of experiments but not another. To understand how to deal with missing values we first constructed an artificial set derived from a high quality published one and eliminated all proteins that have missing values in any of the component protein measurements. From this subset of data with no missing values, we can simulate missing values, knowing what the actual measured values were. To do this we chose a published dataset of different breast cancer cell lines grown in culture, each measured in biological duplicate (Lapek, 2017). We first removed any proteins that have at least one missing value in any of the measured conditions (Fig 2A). Also, we only considered unique peptides that deterministically map to a single protein to avoid circumstances such as shared peptides between different isoforms that may lead to overfitting (i.e. allow the linear model to cheat). The resulting reduced baseline dataset contains comprehensive, quantitative measurements for 6,043 proteins in 82 different conditions with no missing values and no razor peptides. We simulate missing data in this baseline data set by removing some of the values for which we know the ground truth. Our goal is then to impute these values. We started by removing 18 of the 82 different conditions (∼22% data missing). Note that the dataset is log transformed and standardized so that the magnitude of coefficients obtained are more easily interpretable.

**Figure 2.**
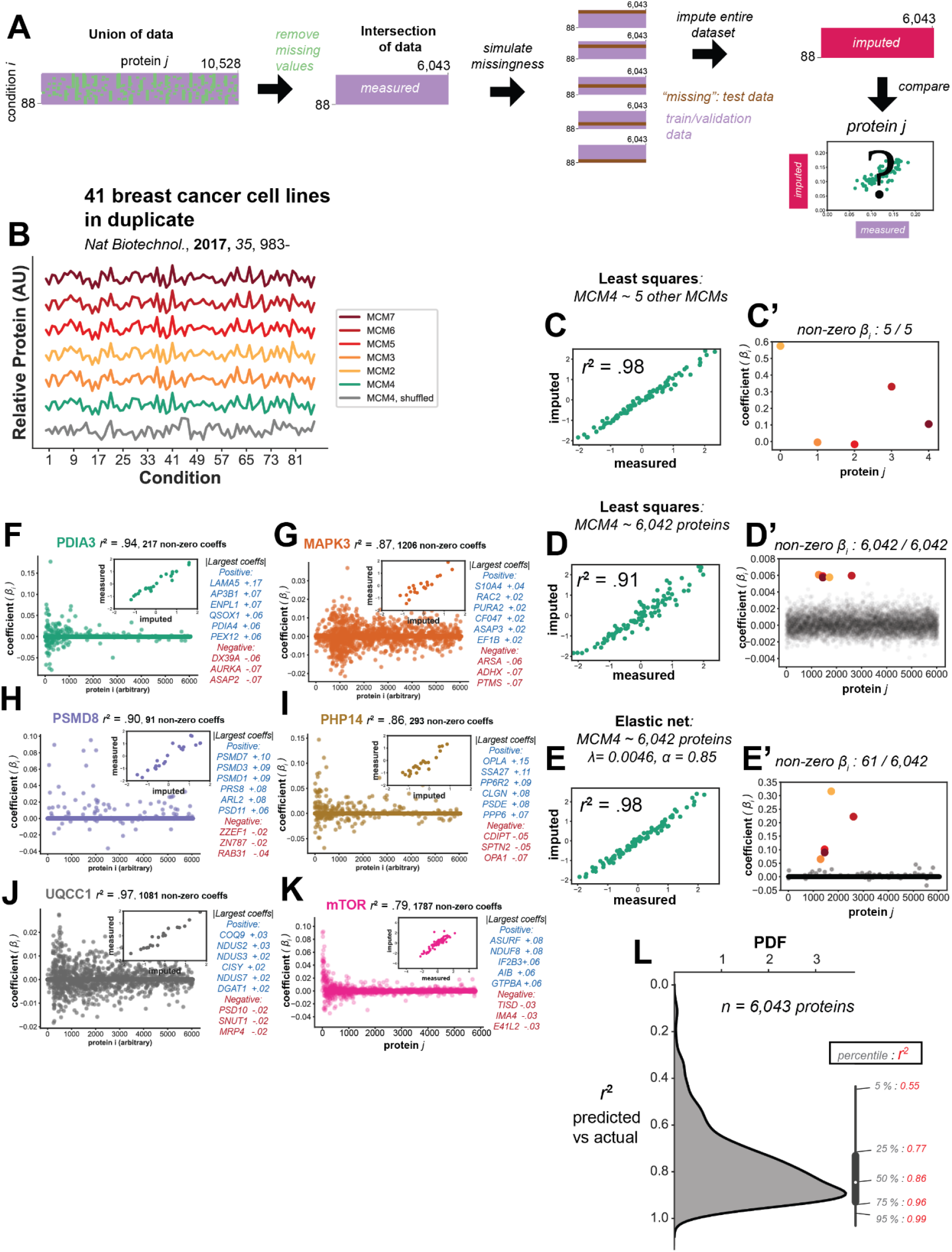
Missing values can be imputed using regularized regression in a multiplexed breast cancer cell line dataset. **A)** All missing values are removed from a sequentially inner merged version of 41 breast cancer cell lines measured in duplicate. The remaining data with no missing values is then used to simulate missingness. “Missing values” are used as a ground truth in comparing imputed data. The dataset is partitioned into a test set that is never used during training, and training and validation sets that are used during cross-validation to select hyperparameters. Missing values are inferred using an elastic net that does a 100 x 20 grid search to select the optimal sparsity and shrinkage parameters. The best model is then selected and compared to the actual values. **B)** Example of the relative abundance of MCM protein complex members in each cell line (colored) and compared to MCM4 data shuffled without replacement (grey). Note the extremely high correlation of all 6 protein complex members (pairwise Pearson *r*^*2*^ range from 0.96 to 0.98 for different complex members). **C) and C’)** Ordinary least squares imputed protein abundance for MCM4 when modeled using the other five members of the complex alone, and the magnitude of the coefficients from the linear model. **D) and D’)** Ordinary least squares imputed protein abundance for MCM4 when modeled using measurements for all 6,402 other proteins in the dataset. A worse, but still good prediction is obtained, but the magnitude of the coefficients is now statistically uninterpretable: all 6,042 proteins used in the model have coefficients with a non-zero magnitude. **E) and E’)** The best elastic net model (from a 200 x 20 grid search) for MCM4 using all 6,042 protein measurements as inputs. Note that not only is an improved prediction obtained, but it is done using a much simpler model. The elastic net used performs variable selection: only 61 / 6,042 proteins in the dataset have a non-zero coefficient. **F)** Best elastic net model coefficients and prediction for PDIA3. **G)** Best elastic net model coefficients and prediction for MAPK3. **H)** Best elastic net model coefficients and prediction for PSMD8. **I)** Best elastic net model coefficients and prediction for PHP14. **J)** Best elastic net model coefficients and prediction for UQCC1. **K)** Best elastic net model coefficients and prediction for MTOR. **L)** Distribution of predicted vs actual Pearson *r*^*2*^ values for all 6,043 proteins in the dataset was plotted using kernel density estimation (KDE) with a gaussian kernel and bandwidth of 0.1.

To highlight the advantageous features of regularized regression, we first compare it to ordinary least squares regression for imputing a member of the MCM protein complex (highly conserved pre-initiation complex for DNA replication found in all eukaryotes) using the other members. The relative abundance measurements in each cell line of the six members of the complex (MCM2, MCM3, MCM4, MCM5, MCM6 and MCM7) as well as randomized data are shown (Fig 2B). Pairwise Pearson correlations are extremely strong (0.96 - 0.98) among members of this discrete complex. Not surprisingly, when we model the abundance of MCM4 as a function of the other members of the complex with ordinary least squares regression, we obtain exceptionally strong predictions with a Pearson *r*^*2*^ of 0.98 when compared to the actual measurements (Fig 2C). All five coefficients from the model are non-zero. Nevertheless, the Pearson coefficients from several of the MCM proteins are disproportionately large because the abundances are essentially co-linear (Fig 2C’). All five coefficients from the model are non-zero. Next, using cross-validation we modeled MCM4 abundance with least squares using all 6,042 protein measurements in the dataset. Though the fit we obtain is still good with an r^2^ of 0.91 (Fig 2D), it is now significantly poorer than we find using only the other 5 members of the MCM complex alone (Fig 2C). Furthermore, each of the 6,042 proteins used in the model has a non-zero coefficient. It is extremely unlikely all 6,042 proteins have some predictive relationship with MCM4. As these proteins will have noise in their measurements, adding these additional measurements weakens the prediction. Thus, the least squares regression fits the noise and therefore actually reduces the correlation over that achieved by using the five MCM proteins alone. A regularized elastic net model can address this problem by removing spurious coefficients that do not correlate with the protein of interest.

To construct an elastic net regularized regression model (which contains 6,042 coefficients that may or may not take on a non-zero value) for all other protein abundances (eqn 1), we perform a large 30 by 30 grid-search over different values of *λ* and *α* to create 900 models with differing amounts of shrinkage and sparsity. For some proteins, only a few proteins in the model will be necessary to strongly predict the abundance, whereas in more complex cases small but real covariation from a large number of proteins may be needed to obtain an accurate prediction. The best elastic net model is selected with cross-validation using a training and validation set. For example, in the case of MCM4, a model with moderate sparsity and shrinkage can be chosen (Fig 2E). Despite having only 61 proteins in the model with non-zero coefficients (Fig 2E’), it yields a prediction that is as accurate as the original least squares, where we provided “the answer” from our previous biological knowledge (Fig 2C). Notably, without any pre-specification, the elastic net correctly assigns the five largest non-zero coefficients to other members of the MCM complex (Fig 2E’). Of note but perhaps not surprising almost all the other non-zero coefficients appear to be proteins involved in DNA synthesis and repair.

### Results of the applicability of Regularized Regression on unmanipulated data sets

We examined the actual coefficients used in elastic net inference for several proteins from the breast cancer cell line model data set (Fig. 2F – 2K). Due to the large range of sparsity and shrinkage that is obtained during the grid search of 900 different elastic net models, some protein models will have imputed values that use only a few predictors, where the coefficients are of large magnitude, whereas others will produce models many coefficients each with a smaller magnitude.

The proteasomal regulatory subunit, PSMD8 (Fig 2H) serves as another example of a defined protein complex, like the MCM complex. The model selected by the elastic net is sparse and contains almost exclusively positive coefficients that correspond to other proteins that form a protein-protein interaction complex with PSMD8. This is reminiscent of work done by many other labs who have had great success in identifying protein-protein interactions by clustering data with sources of biological variation (Pan, 2018). However, this clearly is not the case for all proteins. In the cases of UQCC1 (Fig 2J), MAPK3 (Fig 2G) and mTOR (Fig 2K), there are a comparatively large number of non-zero coefficients (12, 13, and 19 fold respectively), but despite the absence of a few extremely correlated proteins, a comparably strong prediction is obtained. The number of significant predictors may not be surprising, since the mTOR complex has a key role as a nutrient sensor, monitoring many pathways in the cell. Nutrient levels vary up and down and we would expect that some proteins show a corresponding sign of covariation with mTOR. We recognize that although the derived correlations are useful for specifying the level on an unmeasured peptide or protein, the nature of these predictors can be useful in uncovering novel biology, as discussed below. For now, our goal is simply to demonstrate our ability to *impute missing values of protein abundance* without any claims of biological relevance. Finally, we find intermediate levels of coefficient sparseness chosen by the elastic net as the best model for PDIA3 (2F), a chaperone protein in the endoplasmic reticulum that catalyzes isomerization of disulfide bonds, and PHP14 (2I), an understudied phosphatase.

The distribution of Pearson *r*^2^ values obtained from the elastic net prediction and the actual value for all 6,043 proteins obtained from the dataset is shown in (Fig. 2L). Surprisingly, we observe a very high median *r*^2^ of 0.86. The method appears to be robust: the 5^th^ percentile still has an *r*^2^ of 0.55, and 25 percent of the predictions have *r*^2^ that are 0.96 or higher.

### Out-of-fold validation against a permutation null and robustness to missingness

We next established “gold-standard” performance on TMT datasets using an out-of-fold (OOF) framework that scores predictions only on values originally observed in the data and contrasts them to a permutation null built by permuting the response within each fold, preserving the covariance of the actual data matrix used during the imputation. Across random protein subsets and multiple permutations, the OOF R^2^ distribution for real data is cleanly separated from the permuted-null, with medians far to the right of chance under the same design matrix, demonstrating that predictive signal derives from biological covariation rather than model leakage or overfitting (Fig 3A).

**Figure 3.**
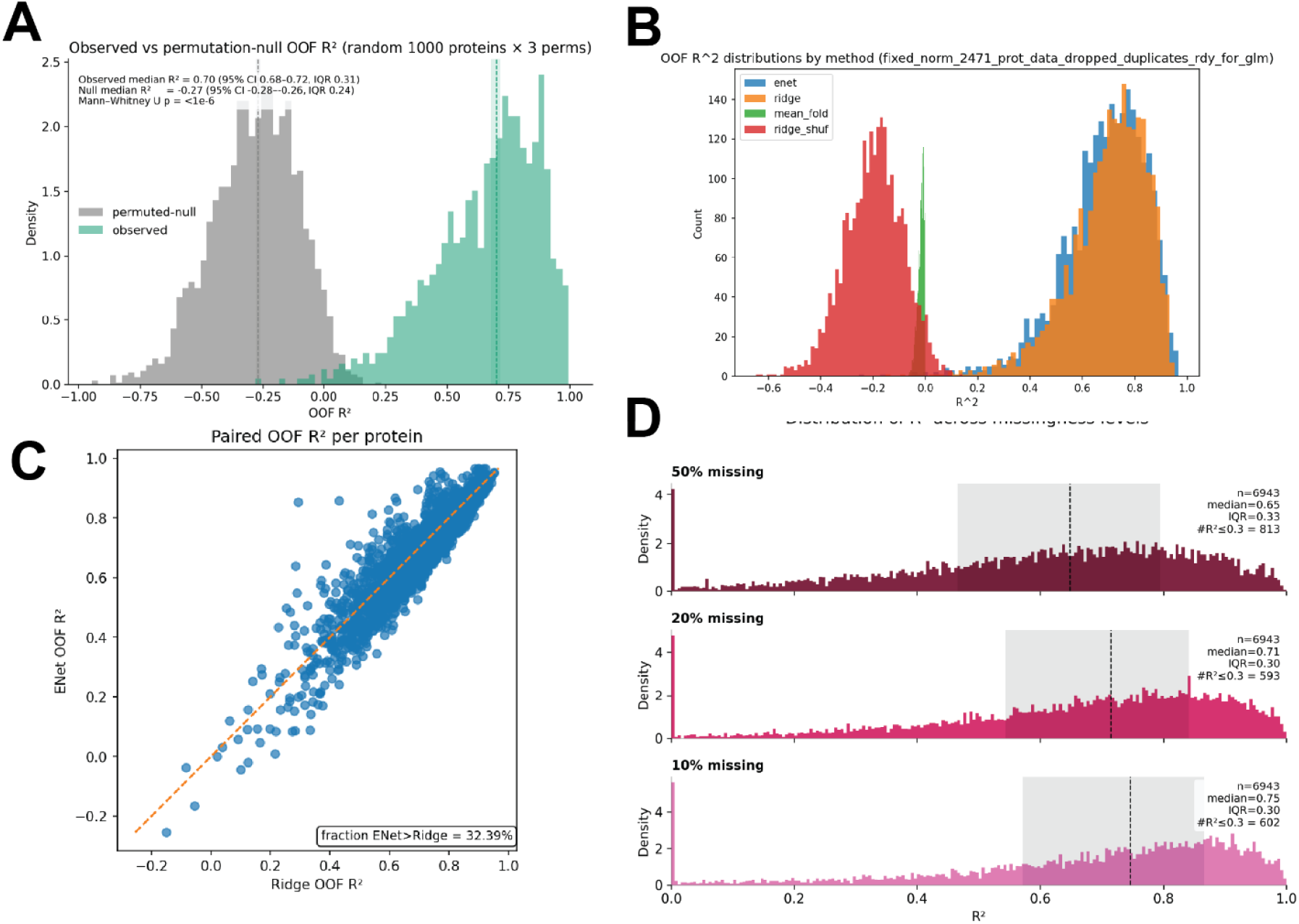
Performance metrics on TMT datasets. **A)** Density overlays of OOF R^2^ for real data (green) versus a permutation null (gray) built by permuting the response within folds (random subset of proteins x multiple permutations). Vertical dashed lines indicate medians. The strong separation establishes the predictive signal is far above chance under the same data structure. **B)** Histograms of OOF R2 across proteins for elastic net (enet), ridge regression (ridge), mean baseline (mean_fold), and the permutation null (ridge_shuf). **C)** Paired Enet vs Ridge OOF R^2^ per protein where each point is one protein and the dashed line is identity. Boxed insert annotation reports the fraction of proteins where Enet > Ridge, which is 32.39%. The tight near diagonal cloud shows the two linaer models perform similarly on most proteins, with Enet sometimes trading a small amount of R^2^ for sparsity and interpretability. **D)** Distributions of OOF R^2^ when masking 10%, 20% or 50% of values as held-out “truth” to assess robustness to various amounts of missingness. Dashed vertical lines mark medians and grey boxes highlight the interquartile range. Predictive quality degrades smoothly with increased missingness, yet a large fraction of proteins remain well predicted with 50% of data missing.

Method comparisons show consistent gains over naïve baselines. Histograms of per-protein OOF R^2^ place elastic net and ridge well above the train-fold mean baseline (sometimes used in label free studies) and far above the permutation null. In paired analyses, elastic net and ridge lie close to the identity line for most proteins, indicating similar accuracy on the majority; nevertheless, elastic net exceeds ridge for roughly one third of proteins (boxed inset, ∼32.4%), reflecting cases where sparse variable selection improves generalization while retaining interpretability.

Finally, we tested robustness to increasing sparsity by masking different fractions of entries as held out truth before fitting. Predictive quality degrades smoothly as missingness rises from 10% to 50%, yet a large fraction of proteins remain well predicted even at 50% masked. Thus, performance scales gracefully with the degree of missingness and supporting the use of the model across studies with widely varying completeness.

### Imputation generalizes across TMT, DIA, label free, and phosphoproteomics

We next asked whether covariation based imputation performs reliably across major proteomic modalities. Using a single analysis workflow, we benchmarked elastic net predictions against the permutation null. In four representative TMT datasets (Fig. 4A–D), the per protein OOF R^2^ distributions lie far to the right of the null with high medians and long high accuracy tails, indicating that much of the proteome is predictable from the remainder of the proteome rather than by chance. The same pattern holds for DIA (Fig. 4E–F) and for a label free study (Fig. 4G), demonstrating that performance is not tied to multiplexing or acquisition strategy. The method also works with phosphopeptide data where an extreme amount of data missingness is often observed (Fig 1E). For a TMT phosphoproteomics of 215 samples in different yeast strains, the OOF R^2^ distribution for phosphopeptides remains well separated from the null (Fig. 4H), despite lower single run abundances and intrinsically sparser sampling.

**Figure 4.**
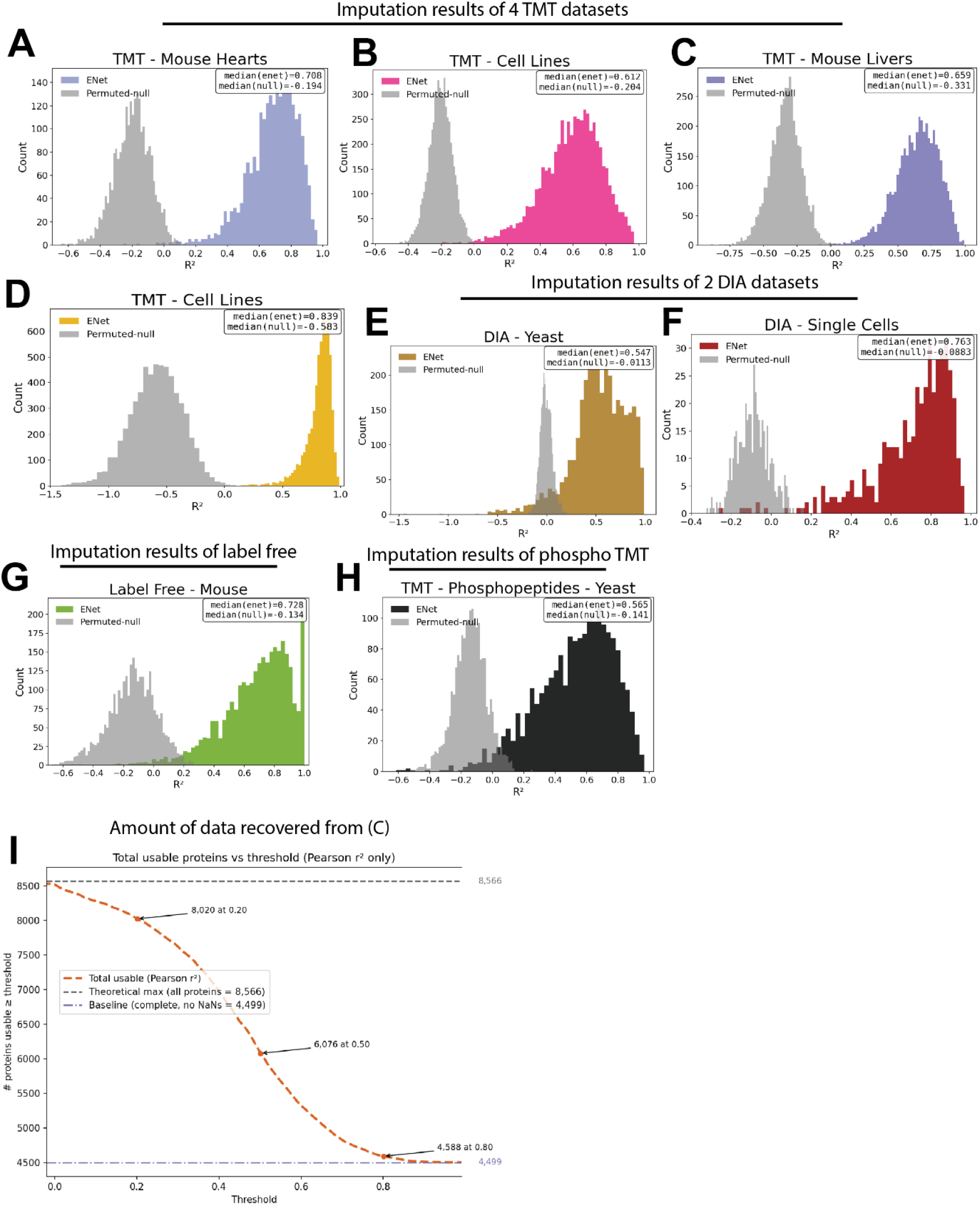
Accurate imputation of missing data across many popular proteomic modalities (TMT, DIA, label free, TMT-phosphoproteomics). All datasets were analyzed identically (log scaled, per feature standardized). Imputation quality is evaluated OOF on originally observed values. The permutation null permutes *y* within each fold to preserve the covariance of *X*. Histograms show the distribution OOF R2 across proteins/sites for ENet on the actual data (colored) versus ENet on permuted null data. **A-D)** Four representative TMT datasets. In every case the ENet distribution is far to the right of the permutation null, with a high median R^2^ and a long high accuracy tail. **E-F)** Two DIA datasets analyzed with the same OOF design. ENet again substantially outperforms the null, demonstrating the approach generalizes beyond multiplexed proteomics. **G)** A label free dataset analyzed with the same OOF design. **H)** TMT phosphopeptides analyzed with the same OOF design. **I)** The number of proteins from 5C that become usable as a function of an imputation quality threshold (here based on Pearson *r* with ground truth OOF targets). The dashed purple line marks the complete case baseline (no missing values). The grey dashed line marks the theoretical maximum. Imputation substantially increases the analyzable set. In this dataset, 6,076 proteins meet *r* ≥ 0.5 versus 4,499 by complete case only, and 8,020 meet *r* ≥ 0.20 (approaching the theoretical maximum of 8,566).

These gains translate into substantially larger analyzable matrices when we gate features by an accuracy threshold. Using the dataset in Fig. 4C as an example, a complete dataset for 6,076 proteins is achieved with *r*^2^ ≥ 0.50 after imputation versus 4,499 under complete case analysis (Fig 4I). With a threshold *r*^2^ ≥ 0.20, a resulting complete dataset of 8,020 proteins is achieved, which approaches the theoretical maximum of 8,566 proteins present in at least one sample (Fig. 4I). Thus, a single, out-of-fold regularized regression procedure generalizes across TMT, DIA, label free, and phosphoproteomics, recovering thousands of otherwise discarded proteins while maintaining stringent, per-feature accuracy control.

### Imputed data preserves global biological structure

On a TMT dataset of ∼7,000 proteins quantified from brown fat in 185 diversity outbred mice, we compared commonly used unsupervised learning approaches on the published data to an entirely imputed dataset with 20% data missing (generated with five 20% OOF, slices). The PCA spectra for measured and imputed matrices are almost identical and strongly diverge from the shuffled-label null (Fig. 5A), indicating that variance is redistributed in the same way after imputation while the null collapses structure into the first component. Component loadings align almost perfectly: imputed vs measured loadings for PC1 and PC2 fall on the identity (r^2^≈1; Fig. 5B), and the mean loading bias (imputed–measured) is ∼0 across the loading range and remains within the null 95% band (Fig. 5C). The distribution of loading differences is sharply centered at 0 and far narrower than the null (Fig. 5D), arguing against strong systematic inflation/deflation of feature weights.

**Figure 5.**
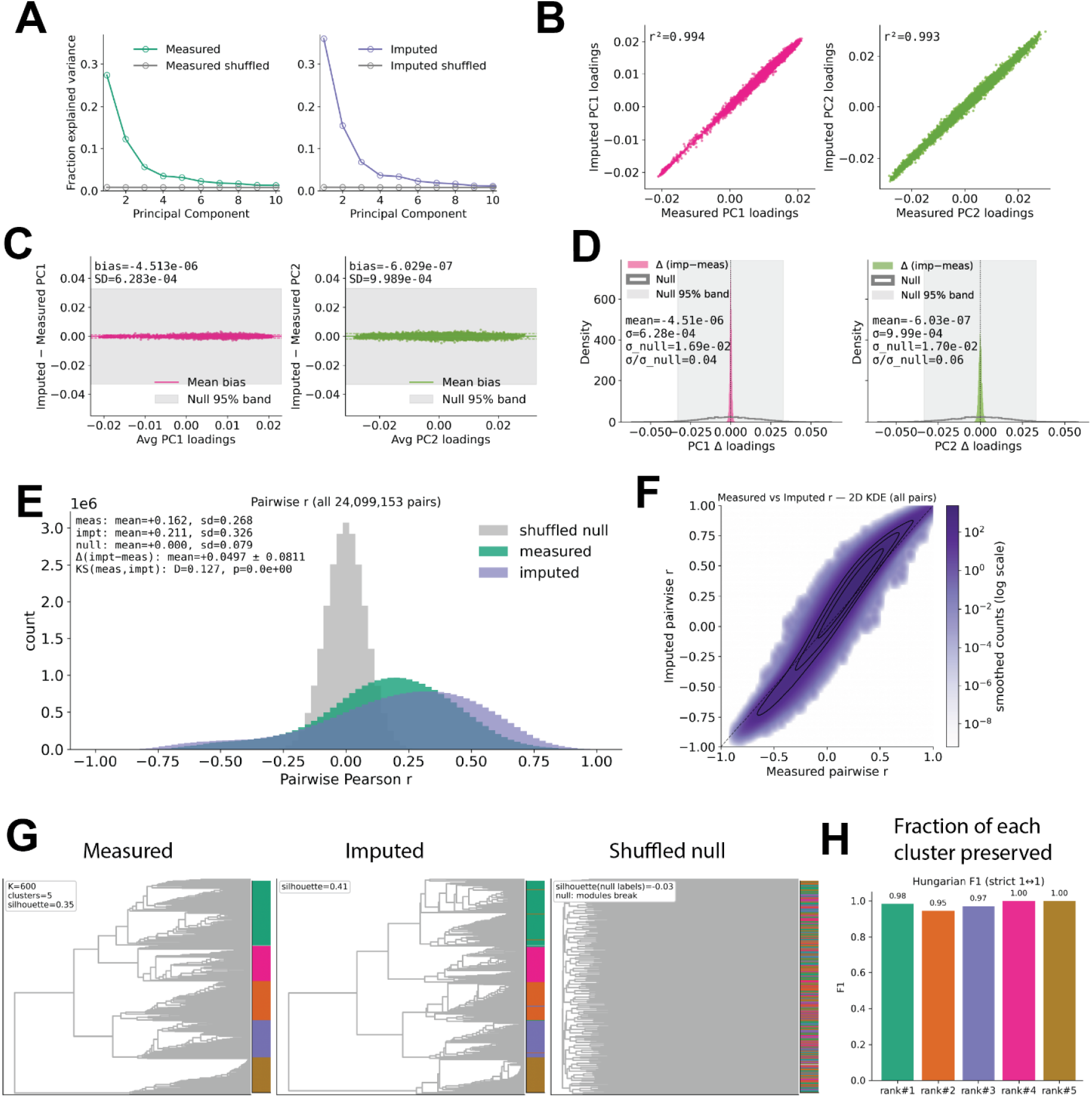
Biological structure of imputed data very closely resembles measured data on a representative TMT dataset. **A)** Fraction of variance explained by the first 10 PCs for measured (left) and imputed (right), each overlaid with their shuffled null spectra. Spectra for measured vs imputed nearly coincide, while shuffling collapses variance into the leading PC and flattens the tail, indicating preserved global covariance after imputation. **B)** Loading scores for PC1 and PC2: imputed vs measured, with sign aligned. Points sit tightly on the identity (r^2^≈1 for both PCs), showing that imputation preserves sample placement along major biological axes. **C)** For PC1 and PC2, mean difference in loadings (imputed – measured) as a function of the average loading. Grey shaded bands are the shuffled null 95% intervals. Mean bias is ∼0 and sits inside the null band, indicating no systematic inflation/deflation of feature weights. **D)** Density of (imputed – measured) loadings for PC1 and PC2 (grey = null). The imputed-measured distributions are sharply centered at 0 and far narrower than the null, confirming minimal distortion. **E)** Distributions of all pairwise Pearson *r* between samples for measured (green), imputed (purple) and shuffled null(grey). **F)** Measured vs imputed pairwise *r* 2D density plot where every sample-sample pair is plotted once (log-scaled density). **G)** Hierarchical clustering at K = 600. The color bar shoes the measured-subset module labels projected onto each dendrogram. In all panels the color stripe encodes measure-subset module identity (which is contiguous along each measured cluster). **H)** Preservation of each protein module in the imputed data. Bars show Hungarian F1 (strict 1 to 1 pairing between measured and imputed clusters). Scores are uniformly high with ≥ 95% of all proteins retained in each module.

At the sample–sample level, measured and imputed pairwise Pearson correlations have nearly overlapping distributions, whereas the shuffled null is centered near zero (Fig. 5E). Some overlap of both the measured and imputed data with the null is expected, as many protein-protein pairs are not biologically correlated, so their true correlations cluster around zero. Plotting imputed versus measured *r* for every pair yields a tight near identity ridge with modest shrinkage around *r* ≈ 0(Fig. 5F). Consistent with this, a direct bias profile (Δ*r* = *r*_imputed_ − *r*_measured_) shows small, systematic deviations: slightly negative for moderately negative pairs and slightly positive for moderately positive pairs, approaching zero at the extremes (Fig. S5). This reflects the expected regularization induced shrinkage that can modestly inflate leading variance components (e.g., PC1), while largely preserving the sign and relative ranking of pairwise relationships.

Finally, hierarchical clustering of proteins identifies the same biological “modules” in the imputed data as those in the measured data. Here, a module simply means a cluster of proteins whose abundance profiles strongly covary across samples. We defined modules by cutting the measured data dendrogram at K = 600 (after removing singletons), then asked whether those same groups remain contiguous when the measured labels are projected onto the imputed dendrogram. We deliberately “over segmented” the measured dendrogram at K = 600 to create small, tight clusters with high within cluster correlation. This gives us a conservative set of coherent groups from the measured data with high biological covariation that are unlikely to arise from noise and can serve as ground truth landmarks for evaluation. Defining these modules only on the measured data (and then projecting the labels onto the imputed tree) avoids circularity: we test whether the same biologically coherent groups reappear after imputation rather than rediscovering them from the imputed matrix. We find the vast proportion of each labeled module stay intact, whereas a shuffled label control loses all information (Fig. 5G). Quantitatively, one-to-one preservation is extremely high: Hungarian matched F1 scores are ≈0.95–1.00 for each of the five largest modules (Fig. 5H), meaning that ≥95% of proteins from each measured module are reassigned to a single, matching module in the imputed data. Thus, the clustering structure that biologists interpret as coregulated protein groups is largely maintained after imputation, enabling downstream pathway or complex level analyses on complete matrices.

## Discussion and Conclusion

Our results demonstrate that large proteomic datasets contain sufficient cross protein covariation to enable accurate statistical inference of missing values. Using elastic net regularized regression, we recovered held out protein abundances with median prediction R^2^ values of 0.5 to 0.8 across a range of acquisition modalities, including TMT, DIA, label free DDA, and TMT phosphoproteomics. These results establish that the quantitative structure of proteomic data, previously considered too heterogeneous for predictive modeling, contains reproducible dependencies that can be systematically exploited to complete sparse matrices.

A key observation from this work is that missingness in proteomics is highly structured and scale dependent. As cohort size grows, the intersection of proteins quantified across all samples rapidly collapses, discarding much of the biologically relevant information. Our approach mitigates this limitation by leveraging information already present in the dataset, without the need for external priors or pathway annotations. Because the method is implemented in an out-of-fold design and evaluated against a permutation null baseline, its reported accuracy directly reflects recoverable biological signal rather than overfitting or leakage.

Methodologically, the elastic net framework provides two advantages over traditional least squares or noise model approaches. First, its joint use of L_1_ and L_2_ penalties encourages parsimonious yet stable models, automatically adjusting complexity to the predictive content of each target protein. Second, the per feature out-of-fold evaluation supplies an internal quality metric that can be used to gate or weight imputed values, enabling users to balance coverage and confidence in downstream analyses. The same workflow generalizes across platforms without any tuning beyond standard normalization, suggesting that underlying covariation patterns are a shared property of biological datasets rather than an artifact of a given mass spectrometry acquisition strategy.

In addition to increasing completeness, imputation preserved essential global structure. Principal component spectra, pairwise correlation distributions, and hierarchical clustering modules derived from imputed data closely matched those from measured values. This indicates that regularized regression can reconstruct missing entries while maintaining relative relationships among samples and proteins, which is an important criterion for downstream multivariate analyses such as PCA, network inference, and pathway enrichment.

Although this framework markedly improves data completeness, several limitations should be acknowledged. Prediction accuracy depends on the number of samples and the intrinsic strength of biological covariation: extremely small or binary contrast datasets will yield weaker performance. As missingness increases beyond ∼50%, uncertainty in the imputed values grows, emphasizing the need to report per feature R^2^ thresholds when using imputed data quantitatively. Moreover, while regression coefficients may highlight known complexes or pathways, they should not be interpreted as causal relationships without additional experimental validation.

Overall, regularized regression provides a practical, modality agnostic solution to the pervasive missing value problem in quantitative proteomics. By reconstructing incomplete measurements while preserving the multivariate geometry of the data, it enables more comprehensive and statistically robust analyses of large-scale proteomic studies. Beyond imputation, the learned coefficients often recapitulate known complexes and pathways and occasionally highlight negative or cross pathway associations that are less well cataloged, offering testable hypotheses about regulation that complement traditional analyses. We view this as a second, distinct payoff of covariation based modeling: it provides interpretable, systems level context for protein relationships that can guide targeted experiments.

---

**Figure S1.**
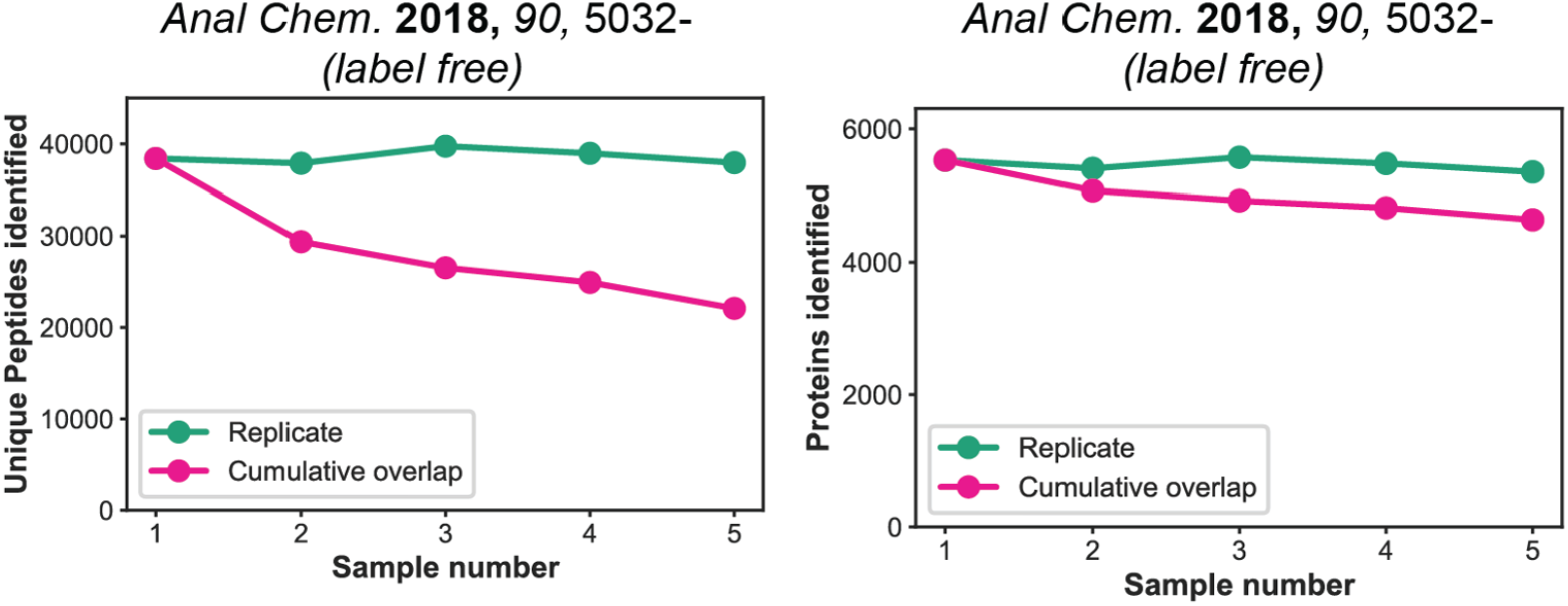
Run to run dropout in label free technical repeats of the same sample. Five back-to-back injections of the *same* tryptic digested HeLa lysate were analyzed on the same mass spectrometer. **Left:** unique peptides identified per run (green) and the cumulative overlap (pink), defined as the intersection of identifications across the first *k* runs. **Right:** the same analysis at the protein level (peptides mapped to proteins). Replicate level counts remain relatively stable, but the cumulative overlap declines with each additional run and shows substantial stochastic missingness even for identical material from a small number of samples. This gap between the union and the intersection motivates imputation when integrating multi-sample datasets.

**Figure S2.**
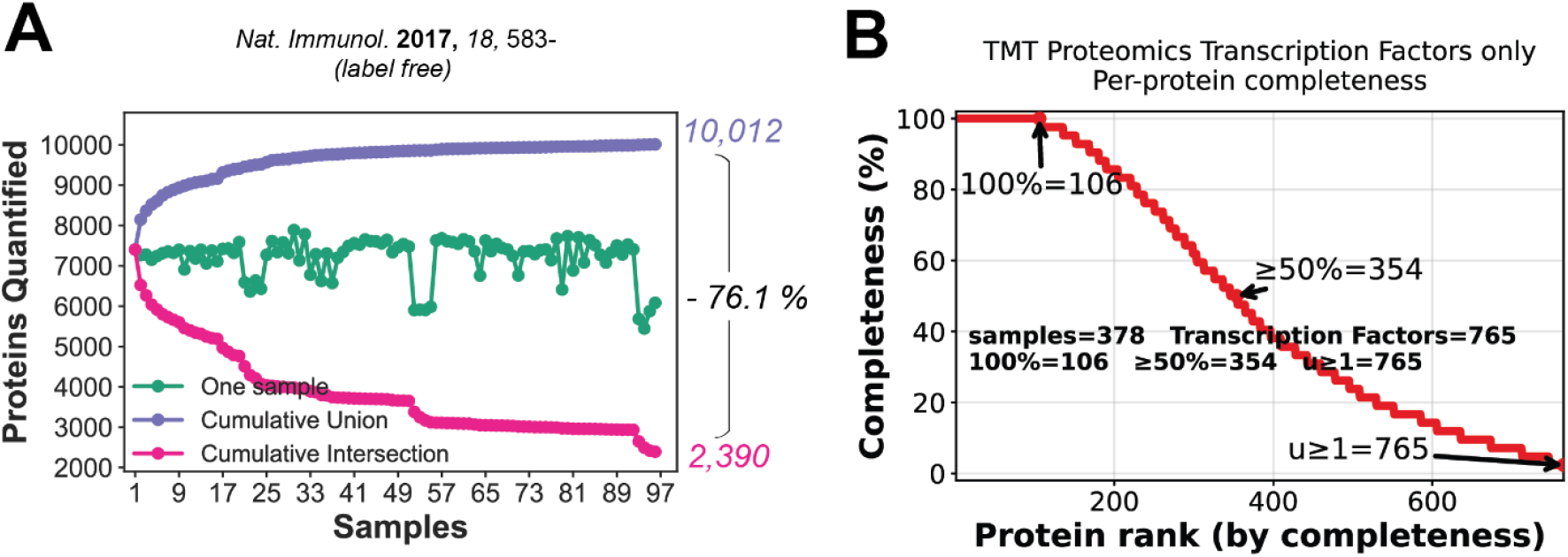
Scale magnifies missing values and disproportionately affects low abundance proteins. **A)** For a label free immune cell atlas of 97 samples of different cell types, the cumulative union reaches 10,012 proteins (purple), whereas the cumulative intersection (pink, proteins quantified in every run) drops to 2,390. Per sample IDs (green) remain roughly constant, but the across sample intersection steadily declines, illustrating run to run dropout and the growth of missingness with cohort size. **B)** Completeness curve for 765 transcription factors (TFs) quantified in at least one of 378 cell lines analyzed with TMT. Only 106 TFs were quantified in 100% of samples, and 354 in ≥ 50%.

**Figure S3.**
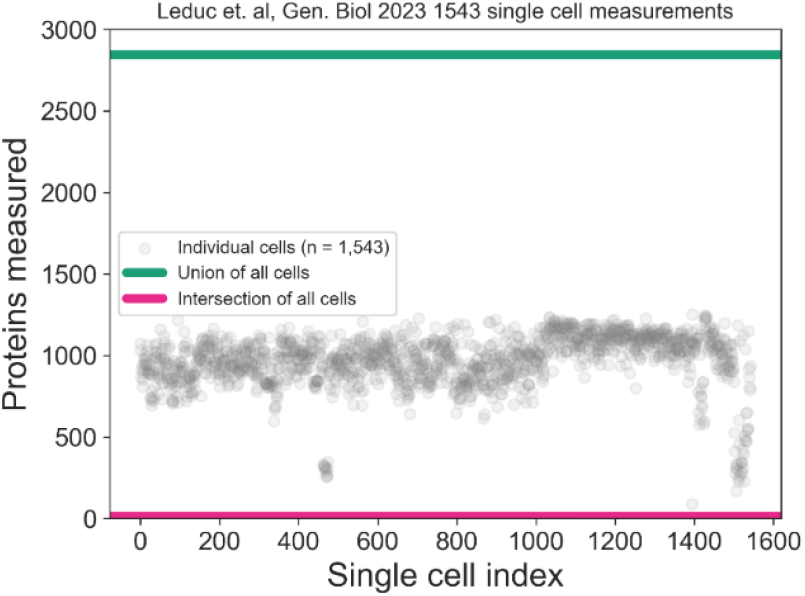
Extreme Sparsity in single cell proteomics. Single cells from Leduc et al., Genome Biology 2023 (*n* = 1,543) are shown as gray points (proteins quantified per cell). The union across all cells (green) totals 2,844 proteins, whereas the intersection (proteins quantified in every cell; pink) is only 13. Despite reasonable per cell depth, the across cell overlap collapses, illustrating the severe run to run/ cell to cell missingness that motivates rigorous imputation for single-cell datasets.

**Figure S4.**
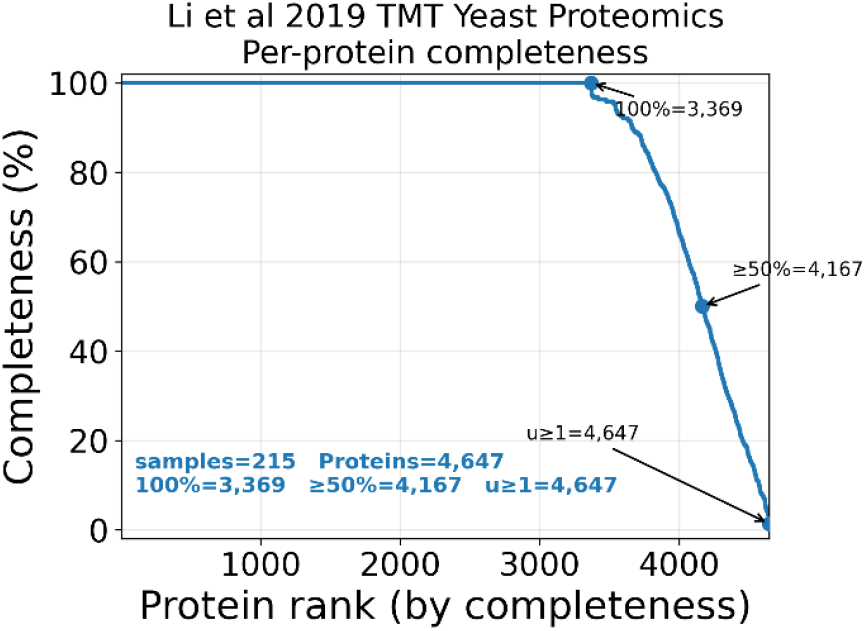
Protein completeness of a large TMT yeast dataset (Li et al., 2019). Per protein completeness curve across 215 samples. Proteins are ranked by completeness (fraction of samples with a quantitative value). Of 4,647 proteins quantified in at least one sample (union), 3,369 are present in 100% of samples and 4,167 are present in ≥50%.

**Figure S5.**
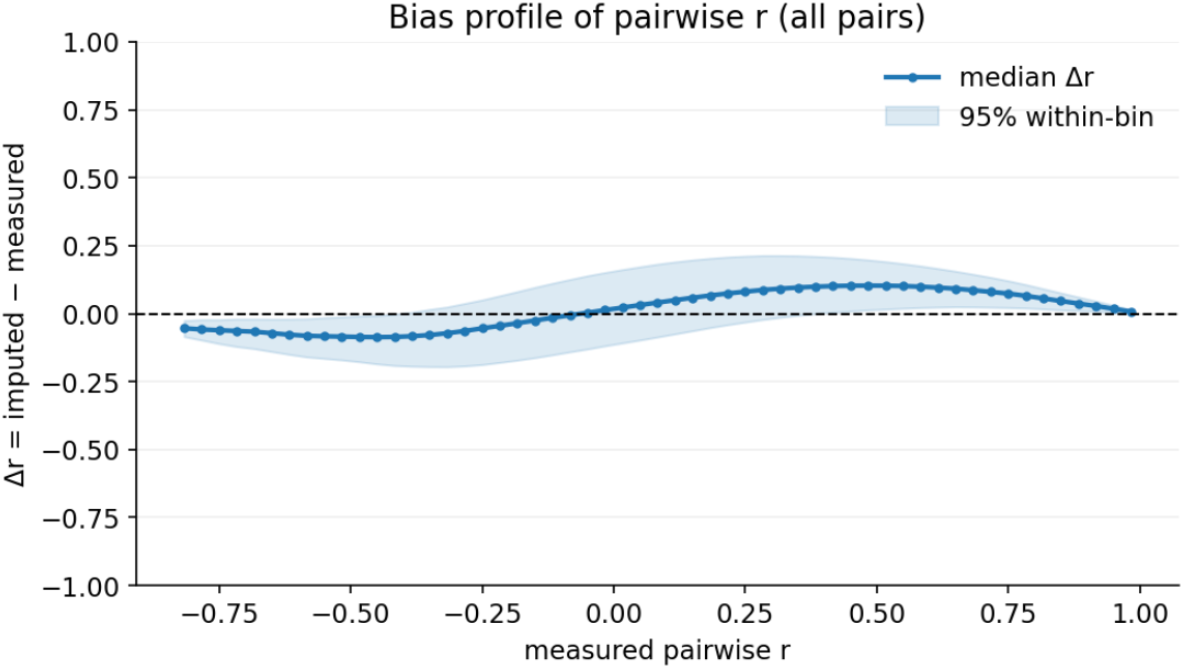
Bias profile of pairwise correlations after imputation. For all protein–protein pairs, we binned by the measured Pearson correlation r(x-axis) and plotted the within bin median Δr = r_imputed_ − r_measured_(line) with the 95% within bin interval (shaded). The profile shows mild negative bias for moderately negative pairs, mild positive bias for moderately positive pairs, and near zero bias at the extremes.

## References

Bludau I, Aebersold R. Proteomic and interactomic insights into the molecular basis of cell functional diversity. Nat Rev Mol Cell Biol. 2020;21(6):327–340.

Chick JM, Munger SC, Simecek P, et al. Defining the consequences of genetic variation on a proteome-wide scale. Nature. 2016;534(7608):500–5.

Giansanti P, Samaras P, Bian Y, et al. Mass spectrometry–based draft of the mouse proteome. Nat Methods. 2022;19(7):803–811.

Golub TR, Slonim DK, Tamayo P, et al. Molecular classification of cancer: class discovery and class prediction by gene expression monitoring. Science. 1999;286(5439):531–7.

Guzman UH, Martinez-Val A, Ye Z, Damoc E, Arrey TN, Pashkova A, Renuse S, Denisov E, Petzoldt J, Peterson AC, Harking F, Østergaard O, Rydbirk R, Aznar S, Stewart H, Xuan Y, Hermanson D, Horning S, Hock C, Makarov A, Zabrouskov V, Olsen JV. Ultra-fast label-free quantification and comprehensive proteome coverage with narrow-window data-independent acquisition. Nat Biotechnol. 2024;42(12):1855–1866.

Gyuricza IG, Chick JM, Keele GR, Deighan AG, Munger SC, Korstanje R, Gygi SP, Churchill GA. Genome-wide transcript and protein analysis highlights the role of protein homeostasis in the aging mouse heart. Genome Res. 2022;32(5):838–852.

Hou W., Ji Z., Ji H, Hicks SC. A systematic evaluation of single-cell RNA-sequencing imputation methods. Genome Biology. 2020;21: 187.

Lapek JD, Greninger P, Morris R, et al. Detection of dysregulated protein-association networks by high-throughput proteomics predicts cancer vulnerabilities. Nat Biotechnol. 2017;35(10):983–989.

Leduc A, Huffman RG, Cantlon J, Khan S, Slavov N. Exploring functional protein covariation across single cells using nPOP. Genome Biol. 2022;23(1):261.

Li J, Paulo JA, Nusinow DP, Huttlin EL, Gygi SP. Investigation of Proteomic and Phosphoproteomic Responses to Signaling Network Perturbations Reveals Functional Pathway Organizations in Yeast. Cell Rep. 2019;29(7):2092-2104.e4.

Lim MY, Paulo JA, Gygi SP. Evaluating false transfer rates from the match-between-runs algorithm with a two-proteome model. J Proteome Res. 2019;18(11):4020–4026.

Linderman G.C., Zhao J., Roulis M., Bielecki P., Nadler B., Kluger Y. Zero-preserving imputation of single-cell RNA-seq data. Nature Communications. 2022;13: 192.

Mitchell DC, Kuljanin M, Li J, Van Vranken JG, Bulloch N, Schweppe DK, Huttlin EL, Gygi SP. A proteome-wide atlas of drug mechanism of action. Nat Biotechnol. 2023;41(6):845–857.

Nusinow DP, Szpyt J, Ghandi M, Rose CM, McDonald ER 3rd, Kalocsay M, Jané-Valbuena J, Gelfand E, Schweppe DK, Jedrychowski M, Golji J, Porter DA, Rejtar T, Wang YK, Kryukov GV, Stegmeier F, Erickson BK, Garraway LA, Sellers WR, Gygi SP. Quantitative proteomics of the Cancer Cell Line Encyclopedia. Cell. 2020;180(2):387-402.e16.

O’brien JJ, Gunawardena HP, Paulo JA, et al. The effects of nonignorable missing data on label-free mass spectrometry proteomics experiments. Ann Appl Stat. 2018;12(4):2075–2095.

Pan J, Meyers RM, Michel BC, et al. Interrogation of Mammalian Protein Complex Structure, Function, and Membership Using Genome-Scale Fitness Screens. Cell Syst. 2018;6(5):555-568.e7.

Rieckmann JC, Geiger R, Hornburg D, et al. Social network architecture of human immune cells unveiled by quantitative proteomics. Nat Immunol. 2017;18(5):583–593.

Sonnett M, Yeung E, Wühr M. Accurate, Sensitive, and Precise Multiplexed Proteomics Using the Complement Reporter Ion Cluster. Anal Chem. 2018;90(8):5032–5039.

Subramanian A, Narayan R, Corsello SM, et al. A Next Generation Connectivity Map: L1000 Platform and the First 1,000,000 Profiles. Cell. 2017;171(6):1437-1452.e17.

Thompson A, Wölmer N, Koncarevic S, et al. TMTpro: Design, Synthesis, and Initial Evaluation of a Proline-Based Isobaric 16-Plex Tandem Mass Tag Reagent Set. Anal Chem. 2019;91(24):15941–15950.

Tibshirani R, Bien J, Friedman J, et al. Strong rules for discarding predictors in lasso-type problems. J R Stat Soc Series B Stat Methodol. 2012;74(2):245–266.

Tran D, Tran B, Nguyen H, Nguyen T. A novel method for single-cell data imputation using subspace regression. Sci Rep. 2022;12(1):2697.

Webb-robertson BJ, Wiberg HK, Matzke MM, et al. Review, evaluation, and discussion of the challenges of missing value imputation for mass spectrometry-based label-free global proteomics. J Proteome Res. 2015;14(5):1993–2001.

Wen ZH, Langsam JL, Zhang L, Shen W, Zhou X. A Bayesian factorization method to recover single-cell RNA sequencing data. Cell Rep Methods. 2021;2(1):100133.

Xiao H, Bozi LHM, Sun Y, Riley CL, Philip VM, Chen M, Li J, Zhang T, Mills EL, Emont MP, Sun W, Reddy A, Garrity R, Long J, Becher T, Vitas LP, Laznik-Bogoslavski D, Ordonez M, Liu X, Chen X, Wang Y, Liu W, Tran N, Liu Y, Zhang Y, Cy AM, White AP, He Y, Deng R, Schöder H, Paulo JA, Jedrychowski MP, Banks AS, Tseng YH, Cohen P, Tsai LT, Rosen ED, Klein S, Chondronikola M, McAllister FE, Van Bruggen N, Huttlin EL, Spiegelman BM, Churchill GA, Gygi SP, Chouchani ET. Architecture of the outbred brown fat proteome defines regulators of metabolic physiology. Cell. 2022;185(24):4654–4673.e28.

Zou H, Hastie T. Regularization and variable selection via the Elastic Net. J. R. Statist. Soc. B. 2005;67(2):301–320.

Zuniga NR, Frost DC, Kuhn K, Shin M, Whitehouse RL, Wei TY, He Y, Dawson SL, Pike I, Bomgarden RD, Gygi SP, Paulo JA. Achieving a 35-plex tandem mass tag reagent set through deuterium incorporation. J Proteome Res. 2024;23(11):5153–5165.

